# Chromosomal-level genome assembly and single-nucleotide polymorphism sites of black-faced spoonbill *Platalea minor*

**DOI:** 10.1101/2024.04.08.588650

**Authors:** Hong Kong Biodiversity Genomics Consortium, Jerome H.L. Hui, Ting Fung Chan, Leo L. Chan, Siu Gin Cheung, Chi Chiu Cheang, James K.H. Fang, Juan Diego Gaitan-Espitia, Stanley C.K. Lau, Yik Hei Sung, Chris K.C. Wong, Kevin Y.L. Yip, Yingying Wei, Wai Lok So, Wenyan Nong, Sean T.S. Law, Paul Crow, Aiko Leong, Liz Rose-Jeffreys, Ho Yin Yip

## Abstract

*Platalea minor*, the black-faced spoonbill (Threskiornithidae) is a wading bird that is confined to coastal areas in East Asia. Due to habitat destruction, it has been classified by The International Union for Conservation of Nature (IUCN) as globally endangered species. Nevertheless, the lack of its genomic resources hinders our understanding of their biology, diversity, as well as carrying out conservation measures based on genetic information or markers. Here, we report the first chromosomal-level genome assembly of *P. minor* using a combination of PacBio SMRT and Omni-C scaffolding technologies. The assembled genome (1.24 Gb) contains 95.33% of the sequences anchored to 31 pseudomolecules. The genome assembly also has high sequence continuity with scaffold length N50 = 53 Mb. A total of 18,780 protein-coding genes were predicted, and high BUSCO score completeness (93.7% of BUSCO metazoa_odb10 genes) was also revealed. A total of 6,155,417 bi-allelic SNPs were also revealed from 13 *P. minor* individuals, accounting for ∼5% of the genome. The resource generated in this study offers the new opportunity for studying the black-faced spoonbill, as well as carrying out conservation measures of this ecologically important spoonbill species.

## Introduction

The black-faced spoonbill *P. minor* (Threskiornithidae) (Figure 1A) is confined to coastal areas in East Asia including Hong Kong, Macau, Taiwan, Vietnam, North Korea, South Korea, and Japan. The natural habitats of the *P. minor* have been disturbed by human activities and industrialization, leading to the decline in bird’s population over the last century (Takano & Henmi 2012; Guo-An et al 2005)[1,2]. With an estimation of more than 6,000 individuals in the world, the International Union for Conservation of Nature (IUCN) has categorised with globally endangered species. Interestingly, a quarter of the population of *P. minor* in the world can be found in Hong Kong, and it is protected under the Wild Animals Protection Ordinance Cap 200 locally. Genetic methods have been performed and attempted to better retain this species with high conservation value (Lee et al 2017; Li et al 2022)[3,4]. Nevertheless, as of to date, a reference genome for this species remained missing.

**Figure 1.**
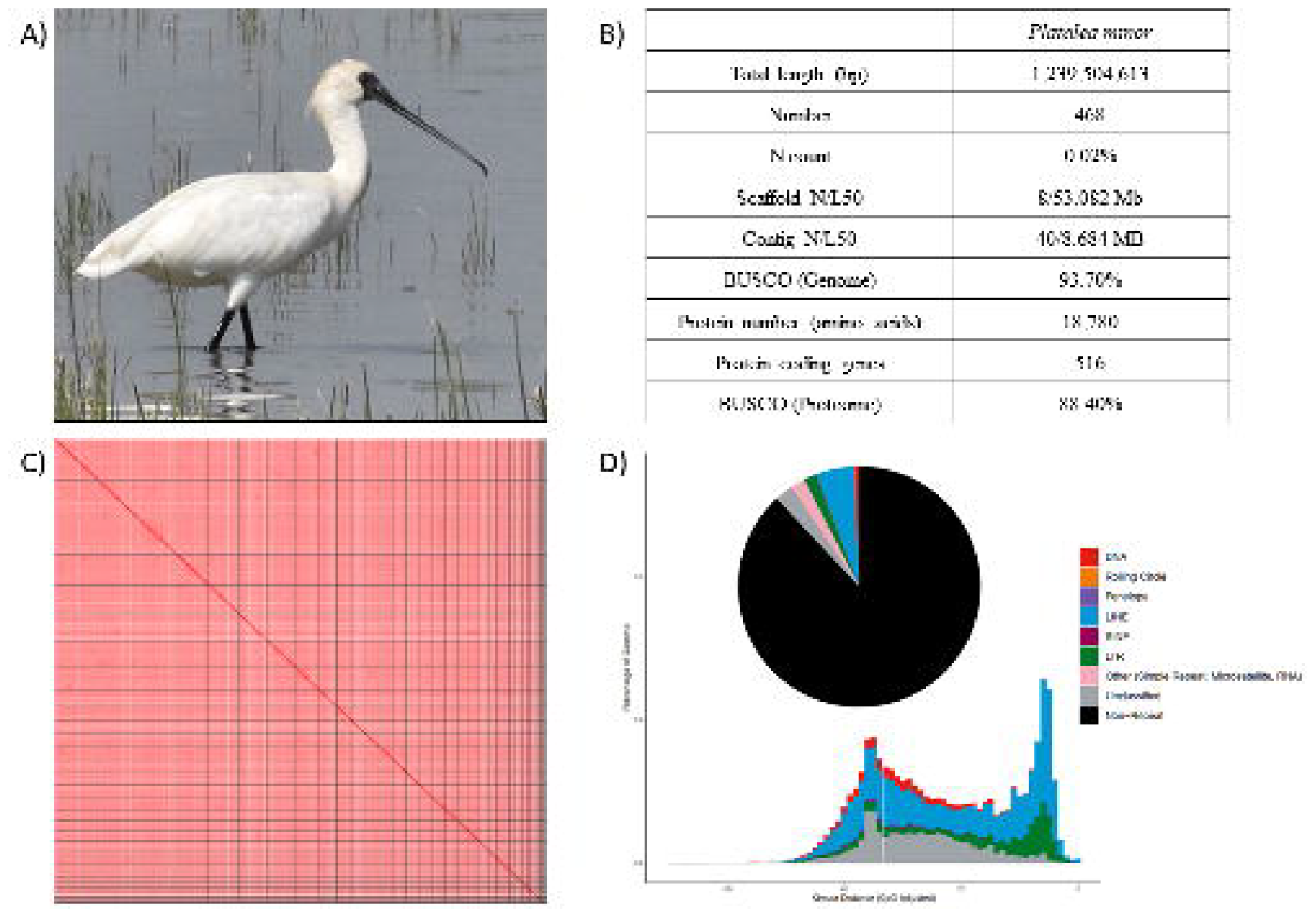
A) Picture of *Platalea minor*; B) Statistics of the genome assembly generated in this study; C) Hi-C contact map of the assembly visualised using Juicebox (v1.11.08); D) Repetitive elements distribution.

## Methods

### Sample collection

Tissue samples of 14 *P. minor* individuals were collected at Tai Po, Hong Kong between February 2015 and February 2020 and subsequently stored in 95% ethanol.

### Isolation of high molecular weight genomic DNA

High molecular weight (HMW) genomic DNA was from a single individual, labelled as “BFS13”. The tissue sample was first ground into powder with liquid nitrogen and subsequently proceeded with the Qiagen MagAttract HMW kit (Qiagen Cat. No. 67563), following the manufacturer’s protocol. The final DNA sample was eluted with 120 μl of elution buffer (PacBio Ref. No. 101-633-500) and was subjected to quality check using the NanoDrop^™^ One/OneC Microvolume UV-Vis Spectrophotometer, Qubit^®^ Fluorometer, and overnight pulse-field gel electrophoresis.

### DNA shearing, PacBio library preparation and sequencing

Approximately 4.4 μg of HMW DNA was processed with DNA shearing through 6 passes of centrifugation in a g-tube (Covaris Part No. 520079) at 2,000 x *g* for 2 min. The sheared DNA was transferred to a 2 ml DNA LoBind^®^ Tube (Eppendorf Cat. No. 022431048) and temporary stored at 4 °C. Overnight pulse-field gel electrophoresis was conducted to assess the fragment size distribution of the sheared DNA. Subsequently, a SMRT bell library was constructed using the SMRTbell® prep kit 3.0 (PacBio Ref. No. 102-141-700), following the manufacturer’s instructions. Briefly, the sheared DNA was processed with DNA repair, followed by polishing and tailing with A-overhang at both ends of each DNA strand. T-overhang SMRTbell adapters were then ligated to the polished ends to form SMRTbell templates, which were purified with SMRTbell^®^ cleanup beads (PacBio Ref. No. 102158-300). The quantity and fragment size of the SMRTbell library were inspected with Qubit^®^ Fluorometer and overnight pulse-field gel electrophoresis, respectively. A nuclease treatment was conducted to remove any non-SMRTbell structures and a subsequent size-selection step with 35% AMPure PB beads was carried out to remove the short fragments. The final preparation of the library was performed using the Sequel^®^ II binding kit 3.2 (PacBio Ref. No. 102-194-100). In brief, Sequel II primer 3.2 and Sequel II DNA polymerase 2.2 were added to anneal and bind to the SMRTbell templates, respectively. An internal control provided by the kit was also added. Finally, the library was loaded on the PacBio Sequel IIe System at an on-plate concentration of 90 pM with the diffusion loading mode. The sequencing was run in 30-hour moves, with a period of 120 min pre-extension. In total, one SMRT cell was used to output HiFi reads and the details of sequencing data are listed in Table 1.

### Omni-C library preparation and sequencing

An Omni-C library was constructed using the Dovetail® Omni-C® Library Preparation Kit (Dovetail Cat. No. 21005), following the manufacturer’s protocol. 80 mg of tissue was ground into powder with liquid nitrogen and was then transferred to 1 mL 1X PBS, followed by crosslinking with formaldehyde and digestion with endonuclease DNase I. An aliquot of 2.5 μL lysate was used for assessing lysate quantification and fragment size distribution using Qubit^®^ Fluorometer and TapeStation D5000 HS Screen Tape, respectively. Then, end polishing, bridge ligation and proximity ligation were carried out in the crosslinked DNA fragments. Subsequently, crosslink reversal was performed, followed by DNA purification and size selection with SPRIselect™ Beads (Beckman Coulter Product No. B23317). The library preparation was continued with end repair and adapter ligation using the Dovetail™ Library Module for Illumina (Dovetail Cat. No. 21004), followed by DNA purification with SPRIselect™ Beads. The DNA fragments were then captured with Streptavidin Beads and Universal and Index PCR Primers from the Dovetail™ Primer Set for Illumina (Dovetail Cat. No. 25005) were added to amplify the DNA library. A final size selection was carried out using SPRIselect™ Beads to retain DNA fragments ranging between 350 bp and 1000 bp. The quantity and fragment size distribution of the library was inspected by the Qubit® Fluorometer and the TapeStation D5000 HS ScreenTape, respectively. The final library was sequenced on an Illumina HiSeq-PE150 platform at Novogene. The details of sequencing data are listed Table 1.

### Genome assembly and gene model prediction

De novo genome assembly was performed using Hifiasm (Cheng et al 2021)[5]. Haplotypic duplications were identified and removed using purge_dups based on the depth of HiFi reads (Guan et al 2020)[6]. Proximity ligation data from the Omni-C library was used to scaffold genome assembly by YaHS (Zhou et al, 2022)[7]. Transposable elements (TEs) were annotated using the automated Earl Grey TE annotation pipeline (version 1.2) as previously described (Baril et al., 2022)[8]. Genome annotation was performed using Braker (v3.0.8) (Hoff et al., 2019)[9] with default parameters. Briefly, the genome was soft-masked using redmask (v0.0.2) (Girgis et al. 2015)[10]. 2,468,534 aves reference protein sequences were downloaded from NCBI as protein hints. A blood RNA-Seq data (SRR6650848) (Cho et al 2019)[11] was also downloaded from NCBI and aligned to the soft-masked genome using hisat2 [12]to generate the bam file. The protein and bam files were used as input to Braker for genome annotation.

### Platalea minor resequencing and single nucleotide polymorphism analysis

Genomic DNA from 13 *P. minor* individuals were isolated using the PureLink™ Genomic DNA Mini Kit (Invitrogen Cat no. K182002), following the manufacturer’s instructions. The quality of DNA samples were assessed with the NanoDrop^™^ One/OneC Microvolume UV-Vis Spectrophotometer and 1% gel electrophoresis and were sent to Novogene for sequencing on an Illumina HiSeq-PE150 platform at approximately 6X coverage. Afterwards, the sequenced raw reads were trimmed by Trimmomatic (v0.39) (Bolger et al., 2014)[13] and cleaned with Kraken 2 (Wood et al., 2019)[14]. The cleaned reads were aligned to large scaffolds (>500 kb, n = 234) that account for 97.1% of the *P. minor* reference genome with BWA-MEM (Li, 2013)[15] using parameters “-t 30 -M -R”. Variant calls were performed using “HaplotypeCaller” and “GenotypeGVCFs” commands from the Genome Analysis Toolkit (GATK, v4.1.2.0) (DePristo et al., 2011)[16]. Hard filtering were employed to filter out single nucleotide polymorphisms (SNPs) with the following criteria: quality by depth (QD) < 2.0, Fisher strand bias (FS) > 60.0, mapping quality (MQ) < 40.0, mapping quality rank sum test (MQRankSum) < -12.5, and read position rank sum test (ReadPosRankSum) < -8.0. The remaining SNPs were further filtered for bi-allelic (“--min-alleles 2 --max-alleles 2) and the heterozygosity and inbreeding coefficient were estimated using VCFtools (v0.1.16) (Danecek et al., 2011)[17].

## Results and discussion

### Genome assembly of P. minor

A total of 25.35 Gb of HiFi bases were generated with an average HiFi read length of 9,365 bp with 20X data coverage (Table 1). After scaffolding with 77.79 Gb Omni-C sequencing data, the assembled genome size was resulted in 1.24 Gb, with 468 scaffolds and a scaffold N50 of 53 Mb in 8 scaffolds (Table 1 and 2; Figure 1B and 1C). The genome size is comparable to the other bird species in the family Threskiornithidae, which have genome sizes around 1.0-1.3 Gb, according to the data available in the NCBI Genbank, such as *Theristicus caerulescens* (1.20 Gb, GCA_020745775.1), *Nipponia nippon* (1.31 Gb, GCA_035839065.1) and *Mesembrinibis cayennensis* (1.19 Gb, GCA_013399675.1). The genome completeness was estimated by BUSCO with a value of 93.7 % (metazoa_odb10) (Table 2; Figure 1B). The GC content was 42.98%. A total of 14,673 gene models were generated with 18,780 predicted protein-coding genes, having a mean coding sequence length of 516 amino acids, and complete protein BUSCO value was 88.4% (Table 2).

### Repeat content

A total repeat content of 11.94% was found in the genome, which contained a lower level of repeat elements, similar to other avian genomes (Zhang et al 2014)[18], with 2.49% unclassified elements. Of the remaining repeats, LINE is the most abundant (5.10%), followed by LTR (1.62%), whereas DNA, SINE, Penelope and rolling circle are only present in low proportions (DNA: 0.63%, SINE: 0.09%, Penelope: 0.06%, rolling circle: 0.02%). A complete catalogue of the repeat content of the genome can be found in Table 4 and Figure 1D.

### Single nucleotide polymorphism sites (SNPs)

A total of 6,155,417 bi-allelic SNPs were called from 13 *P. minor* individuals, accounting for ∼0.5% of the genome. The mean observed heterozygosity was 0.145%, which is comparable to 0.109% from a previous study of 11 black-faced spoonbill samples (Li et al 2022)[4] (Table 5). Signals of inbreeding was observed among the samples, with inbreeding coefficient (*F*_IS_) ranging from 0.331 to 0.720 (Table 5).

### Conclusion and reuse potential

This study presents the first chromosomal-level genome assembly and single-nucleotide polymorphism sites of black-faced spoonbill *Platalea minor*, which is a useful and precious resource for further population genomic studies of spoonbills in light of understanding species numbers and conservation.

### Data validation and quality control

During DNA extraction and PacBio library preparation, the samples were subjected to quality control with NanoDrop^™^ One/OneC Microvolume UV-Vis Spectrophotometer, Qubit^®^ Fluorometer, and overnight pulse-field gel electrophoresis. The Omni-C library was inspected by Qubit^®^ Fluorometer and TapeStation D5000 HS ScreenTape.

Regarding the genome assembly, the Hifiasm output was blast to the NT database and the resultant output was used as the input for BlobTools (v1.1.1) (Laetsch & Blaxter 2017)[19]. Scaffolds that were identified as possible contamination were removed from the assembly manually (Figure 2). A statistical kmer-based approach was applied to estimate the heterozygosity of the assembled genome heterozygosity The repeat content and the corresponding sizes were analysed with k-mer 21 using Jellyfish (Marçais & Kingsford 2011)[20] and GenomeScope (Ranallo-Benavidez et al 2020)[21] (Figure 3; Table 6). Benchmarking Universal Single-Copy Orthologs (BUSCO, v5.5.0) (Manni et al., 2021)[22] was used to assess the completeness of the genome assembly and gene annotation with metazoan dataset (metazoa_odb10). HiC contact maps were generated using Juicer tools (version 1.22.01) (Durand et al. 2016)[23], following the Omni-C manual [24].

**Figure 2.**
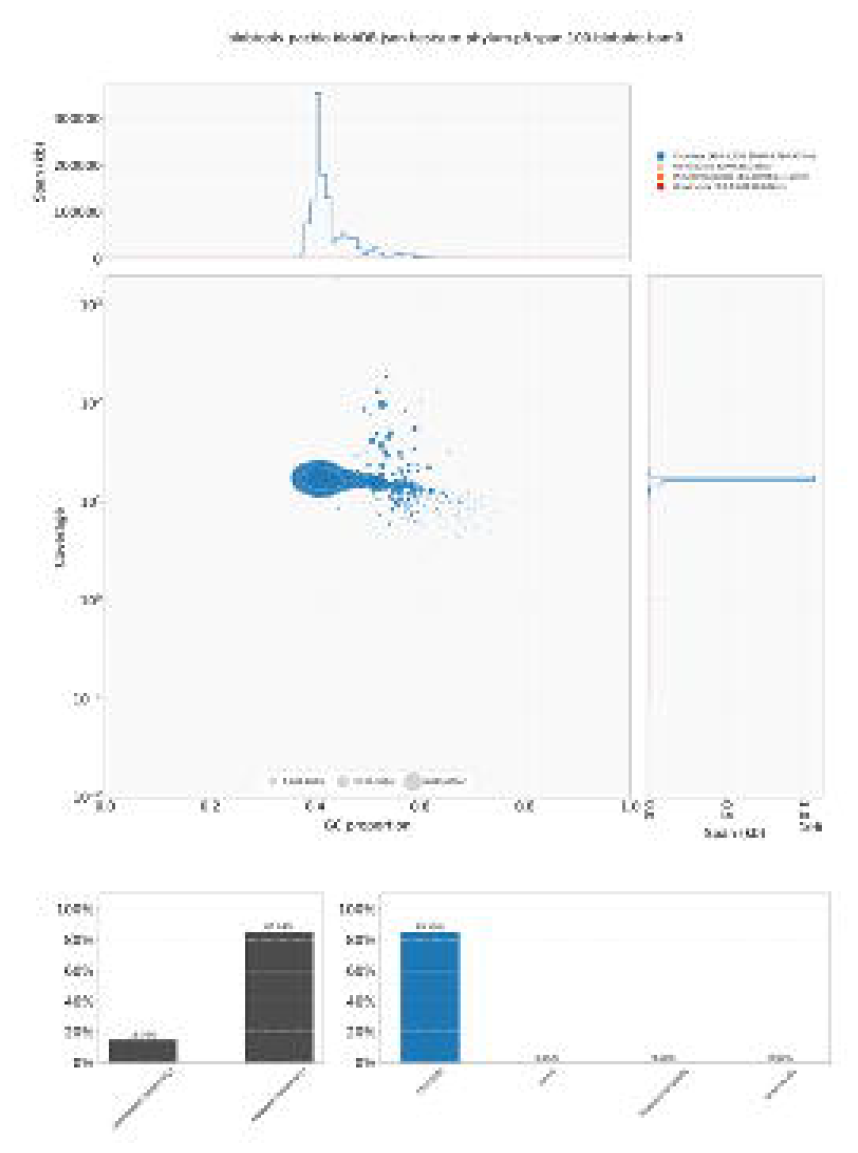
Genome assembly QC and contaminant/cobiont detection. The upper pannel shows the BlobPlot of the assembly, with each circle representing a scaffold with its size scaled according to its scaffold length while the colour of the circle indicates the taxonomic assignment from BLAST similarity search results. The lower pannel reveals the ReadCovPlot of the assembly illustrating the proportion of unmapped and mapped sequences in the BLAST similarity search results on the left. The latter of which is further dissected according to the rank of phylum on the right.

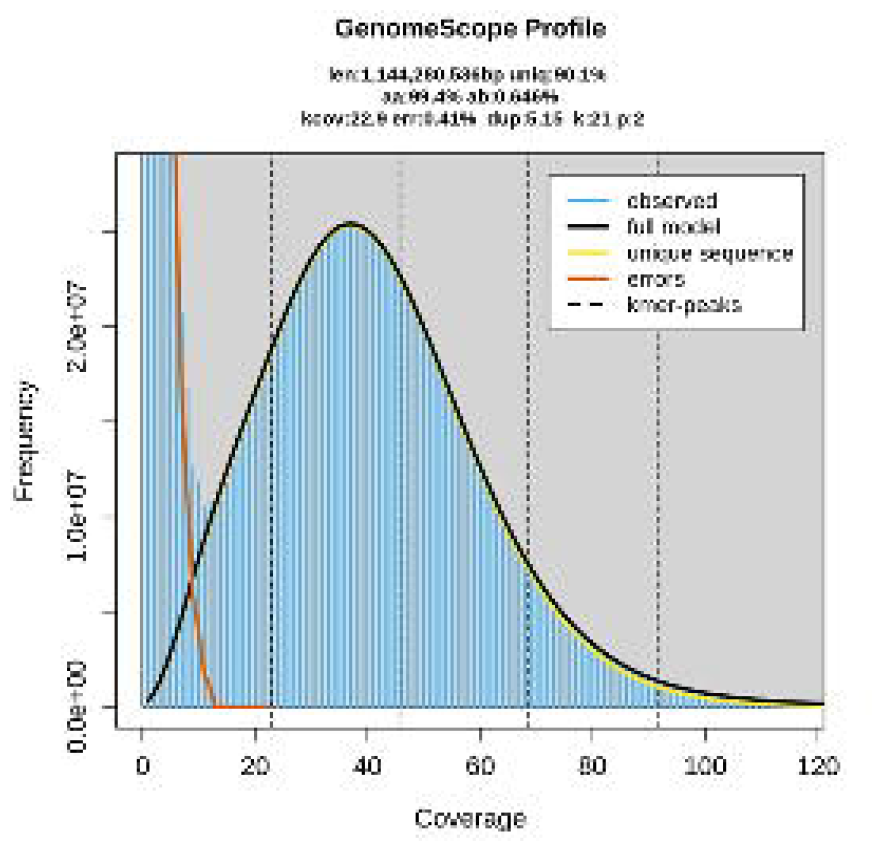

Omni-C reads and PacBio HiFi reads were used to measure assembly completeness and consensus quality (QV) using Merqury (v1.3) (Rhie et al., 2020)[25] with kmer 20, resulting in 95.0738% kmer completeness for the Omni-C data and 59.746 QV scores for the HiFi reads, corresponding to 99.999% accuracy.

## Data availability

The raw reads generated in this study, including Omni-C (SAMN40731791) and PacBio HiFi (SAMN35152374) data, have been deposited in the NCBI database under the BioProject accession number PRJNA973839. The genome, genomic and repeat annotation files have been deposited and are publicly available in figshare (https://figshare.com/s/89f741cde0c1039ce057).

## Funding

This work was funded and supported by the Hong Kong Research Grant Council Collaborative Research Fund (C4015-20EF), CUHK Strategic Seed Funding for Collaborative Research Scheme (3133356) and CUHK Group Research Scheme (3110154).

## Author’s contributions

JHLH, TFC, LLC, SGC, CCC, JKHF, JDG, SCKL, YHS, CKCW, KYLY and YW conceived and supervised the study; WLS carried out DNA extraction, library preparation and sequencing; WN performed genome assembly and gene model prediction; STSL carried out the SNPs calling and Fst calculations; PC, AL, LRJ and HYY collected and maintained the samples. All authors approved the final version of the manuscript.

## Competing interest

The authors declare that they do not have competing interests.

## Table and figure legends

Table 1. Summary of sequencing data

Table 2. Genome statistics

Table 3. Scaffold information with length larger than 1Mb

Table 4. Summary of repetitive elements analysis

Table 5. Number of SNPs and statistics of heterozygosity and inbreeding coefficient of 13 Platalea minor individuals

Table 6. Summary of the GenomeScope statistics (k=21)

## Notes

### Competing Interest Statement

The authors have declared no competing interest.

